# NGBO: Introducing -omics metadata to biobanking ontology

**DOI:** 10.1101/2023.05.09.539725

**Authors:** Dalia Alghamdi, Damion M. Dooley, Mannar Samman, William W.L. Hsiao

## Abstract

**Background:** With improvements in high throughput sequencing technologies and the constant generation of large biomedical datasets, biobanks increasingly take on the role of managing and delivering not just specimens but also data. However, re-using data from different biobanks is challenged by incompatible data representations. Contextual data describing biobank digital resources often contain unstructured textual information incompatible with computational processes such as automated data discovery and integration. Therefore, a consistent and comprehensive contextual data framework is needed to increase discovery, reusability, and integrability across data sources.

**Methods:** Based on available genomics standards (e.g., Minimum information about a microarray experiment (MIAME)), the College of American Pathologists (CAP) laboratory accreditation requirements, and the Open Biological and Biomedical Ontologies Foundry principles, we developed the Next Generation Biobanking Ontology (NGBO). In addition, we created new terms and re-used concepts from the Ontology for Biomedical Investigations (OBI) and the Ontology for Biobanking (OBIB) to build NGBO.

**Results:** The Next Generation Biobanking Ontology https://www.ebi.ac.uk/ols4/ontologies/ngbo is an open application ontology representing omics contextual data, licensed under the Apache License 2.0. The ontology focuses on capturing information about three main activities: wet bench analysis used to generate omics data, bioinformatics analysis used to process and interpret data, and data management. In this paper, we demonstrated the use of the NGBO to add semantic statements to real-life use cases and query data previously stored in unstructured textual format.

## INTRODUCTION

In the last decade, the application of different omic studies (e.g., genomics and transcriptomics) that aimed at understanding a particular problem in human disease has been greatly successful [1]. These studies generate extensive data, which, with careful integration under suitable mathematical frameworks and advanced technologies such as ML, can help solve broader research questions about basic, applied, and personalized health challenges [2]. Biological material and data collection have been significant resources in the modern world for genomics research, translational studies, molecular epidemiology, therapeutic target identification, and biomarker and drug discovery [3]. Therefore, it is unsurprising that researchers from the academic community and industry have shown a growing interest in biobanking specimens, genetic materials, and derivative datasets.

Biobanks have evolved from centers collecting only biological materials to organizations gathering various information and data sets, such as genetic profiles and social, clinical, and pathological records [4].Consequently, biobanks increasingly need robust contextual data - including experimental or monitoring details and technical and analytical methods management to handle heterogeneous data for discovery and integration purposes [5].

Many initiatives have been proposed to encourage and recommend biobanking contextual data management standards. For instance, The BBMRI consortium defined a minimum data set for sharing biobank samples (MIABIS) [6]. MIABIS core terminology consists of three components describing biobanks, where sample collections, studies, and donor information are provided in aggregated form.

Heterogeneity issues, however, remain a burden on biobanking institutions, stakeholders, and users despite the scientific community’s considerable effort [7]. Stored data might be incomplete (due to data silos or collected data being insufficient for data quality concerns, for secondary analysis), inaccurate, or missing contextual data. An example of contextual data for a given sequencing data file is the information on the software program used to assess read quality and filter out poor-quality reads. This provenance information on data processing is not part of the currently available minimum information requirements yet it is specifically required by accreditation protocols. To tackle this challenge, we set up minimum information requirements based on The College of American Pathologists’ documentation requirements [8]. Another example is the semantic ambiguity of the available contextual data. For example, the term “specimen source” could be defined as the “body part which the specimen obtained from” or “ biobanking institution which provides the specimen.” Using ontologies, the formal logical representations of terms prevent unnecessary complexity that can lead to mistakes during the data management and retrieval processes [9]. In the ideal case, the contextual data should provide all necessary information to re-analyze available data, on the one hand, and should enable researchers to reuse data in a broader context going beyond individual studies, on the other hand [10].

Data integration from different biobanks is crucial in many situations to enable statistical analysis to achieve sufficient power. For example, collecting a sufficiently large sample for a rare disease can challenge any institution. Through the coordination of several agencies to combine samples, one can achieve the sample size needed to uncover associations between phenotypes and genotypes [11, 12]. However, there are two key challenges: i) specimen/data discovery and ii) the reusability and integration of the existent data sets available in those biobanks. Complete, coherent, and standard descriptions of contextual data maximize the likelihood for the data to be found, re-used, and shared [13].

Andrade et al. has pointed out that semantically rich ontologies provide a promising approach to discovering and integrating data from diverse biobanks [14]. The first attempt at using ontologies for biobanks was in 2014 when the first version of biobanking domain ontology was published, referred to as Ontologized MIABIS (OMIABIS) [15]. It was later extended (based on defined biobank user needs and scenarios) and combined with Biobanking ontology (BO), forming a novel Ontology for biobanking (OBIB) [16]. One of the key advantages of using ontologies is linking data from biobanks to other biological and biomedical repositories using standard identifiers, such as those from Gene Ontology [17]. Hitherto, no ontology provided the coverage needed to capture a broad spectrum of integrating omics digitals assets into biobanks. This possibility will significantly increase the utility of the shared biobank data since it is simple to retrieve relevant data using semantic web technologies such as RDF, SPARQL, and OWL [16].

We present the Next Generation Biobanking Ontology (NGBO) in this paper. An open biobanking ontology created based on the OBO foundry principles, evaluated and accepted by the OBO foundry, accessible through the Ontology Lookup Services, and withthe Saudi Human Genome Project (SHGP) as an active and supporting use case). Here, we provide details about the methodology used to develop, maintain, and evaluate the ontology. Additionally, we outlined the domain covered by NGBO. Finally, we discuss future steps for expanding the ontology.

## METHODS

We approached the development of NGBO by first answering questions about ontology purpose, scope, intended end-users, and usage (see Table 1). Then, we set out to catalog and study the relevant specifications and existing resources. In this process, we reviewed a selection of public repository requirements (such as National Center for Biotechnology Information (NCBI) BioSample); The College of American Pathologists (CAP) laboratory guidelines; and other currently available guidelines (including minimum information about microarray experiments, genome/metagenome sequences, and high-throughput nucleotide sequencing experiments); relevant American College for Medical Genetics and Genomics (ACMG) guidelines; clinical laboratory reports from King Fahad Medical City; and carefully screened a collection of public contextual data submitted accompanying a selection of PubMed publications (2017–2019). For PubMed publications, we applied a snowball sampling method. We started with relevant papers and selected the next round based on titles, abstracts, and references. The process continued until we collected enough data to answer the NGBO specification questions. From this analysis, we constructed a set of specific data fields for the minimum information requirements for reporting wet bench and bioinformatics analysis processes.

**Table 1.** NGBO specification.

| NGBO specification |  |
| --- | --- |
| Purpose | This ontology is developed to represent omics contextual data to support experts in different data discovery and re-use stages. |
| Scope | This ontology includes real-world physical objects such as specimens, computerized objects such as datasets, and planned processes and related specifications. |
| Intended end user | The intended end-users are biobanking organizations, data collectors, omics researchers, and decision-makers. |
| Intended use | This framework will give a comprehensive view of contextual data. NGBO is intended to be used as a common knowledge management framework. |

To scope and delimit the knowledge (i.e. specific data fields and relations) represented by NGBO, we first clarified the need by proposing the use case scenarios listed below through discussions with domain experts at the SHGP and researchers with previous experience in requesting specimen-derived data from repositories. We then prepared a set of competency questions (CQs) expressed in natural language to evaluate the ontology, which was formalized into description logic (DL) queries according to the language used to represent the ontology. The CQs provide a means to verify the satisfiability of the ontology requirements by retrieving the correct query results.

### Use case scenario 1 - Finding interest data

A researcher must collect specimens and associated data to perform a genome-wide association study (GWAS) for a specific rare disease. While funding agencies, grants, publications, and biobanks increasingly support data sharing, learning that data of interest exists can be challenging. Rich contextual data indexed in a searchable format is required. Contextual data should also be assigned globally unique, persistent identifiers.

### Use case scenario 2 - Integrating data with other data

Following use case scenario 1, GWAS requires a large sample size, making it difficult for a single biobank to provide the necessary data. Researchers may need to integrate data from other sources, applications, and workflows. Contextual data must be associated with community-agreed schemas, rich contextual data that follow relevant standards, and controlled vocabularies, keywords, thesauri, or ontologies that follow FAIR principles[18].

### Use case scenario 3 - Optimizing data reuse

For instance, a researcher must reuse data if they require a set of contextual data to answer a specific research question that differs from the contextual data needed in the primary analysis of the generated data. Data must be associated with rich, detailed provenance to optimize reuse and follow community-standard contextual data. Also, the associated contextual data isolated in data silos or free text must be integrated into a computer-accessible unified data layer with an acceptable data usage license.

The first technical step taken to build NGBO was initiating a GitHub repository using the Ontology Development Kit (ODK), a toolkit for building, maintaining, and standardizing biomedical ontologies, which provides a set of standardized, customizable, and automatically executable workflows and packages for maintaining ontology life cycle [19, 20]. The Protégé ontology editor [21] was used to build the ontology in the Web Ontology Language (OWL) format [22]. To enable the integration of external ontologies and guide how terms interrelate, we followed the Open Biological and Biomedical Ontology (OBO) Foundry Principles [17]. We used Basic Formal Ontology (BFO) as a top-level ontology for NGBO development [23].

With careful assessment, we reused pre-existing terms from other appropriately maintained ontologies when possible to prevent multiple representations of the same entity. These ontologies need to be annotated with proper PURLs, labels, and textual and logical definitions. For example, we directly imported terms from OBI that describe entities involved in biomedical investigations, such as but not limited to material entities, processes, and protocols, through OWL import. Additionally, from the ontology for biobanking [24], we imported terms to describe specimens, donors, and specimen management [24]. Information artifact ontology [25] is another relevant ontology used as a mid-level ontology and includes terms describing identifiers, software solutions, and documents.

Because NGBO is an application ontology, we submit new terms introduced by NGBO to the appropriate OBO Foundry domain ontology when possible. Input from a greater OBO foundry community was solicited during the International Conference on Biomedical Ontologies 2019, where NGBO was first presented. Inputs from use case providers, SHGP, and all comments are incorporated as appropriate.

## RESULTS

NGBO is an open-source application ontology written in OWL, Licensed under the Creative Commons Attribution 4.0 International Public License. Available at a public repository where ontology releases and term requests are maintained, accessed by the Ontology Lookup Service at https://www.ebi.ac.uk/ols4/ontologies/ngbo. The first release, published in Jan 2023, contains 1643 classes; 192 are NGBO-defined, 30 individuals, 32 object properties, and 1942 logical axioms and relations.

The “is_a” relationship (blue arrows in Figures 1-4) represents a relationship that expresses an inheritance between parent and child. Apart from the “is_a” relation, which is commonly used in all domains, NGBO also includes domain-specific relations, such as “execute,” “trace_from,” and “has_specified_input.” The majority of relations were imported from the Relations Ontology to improve interoperability with other ontologies [21]. These relations reflect the domain knowledge structure, representing the semantic connections between the ontology concepts.

**Figure 1:**
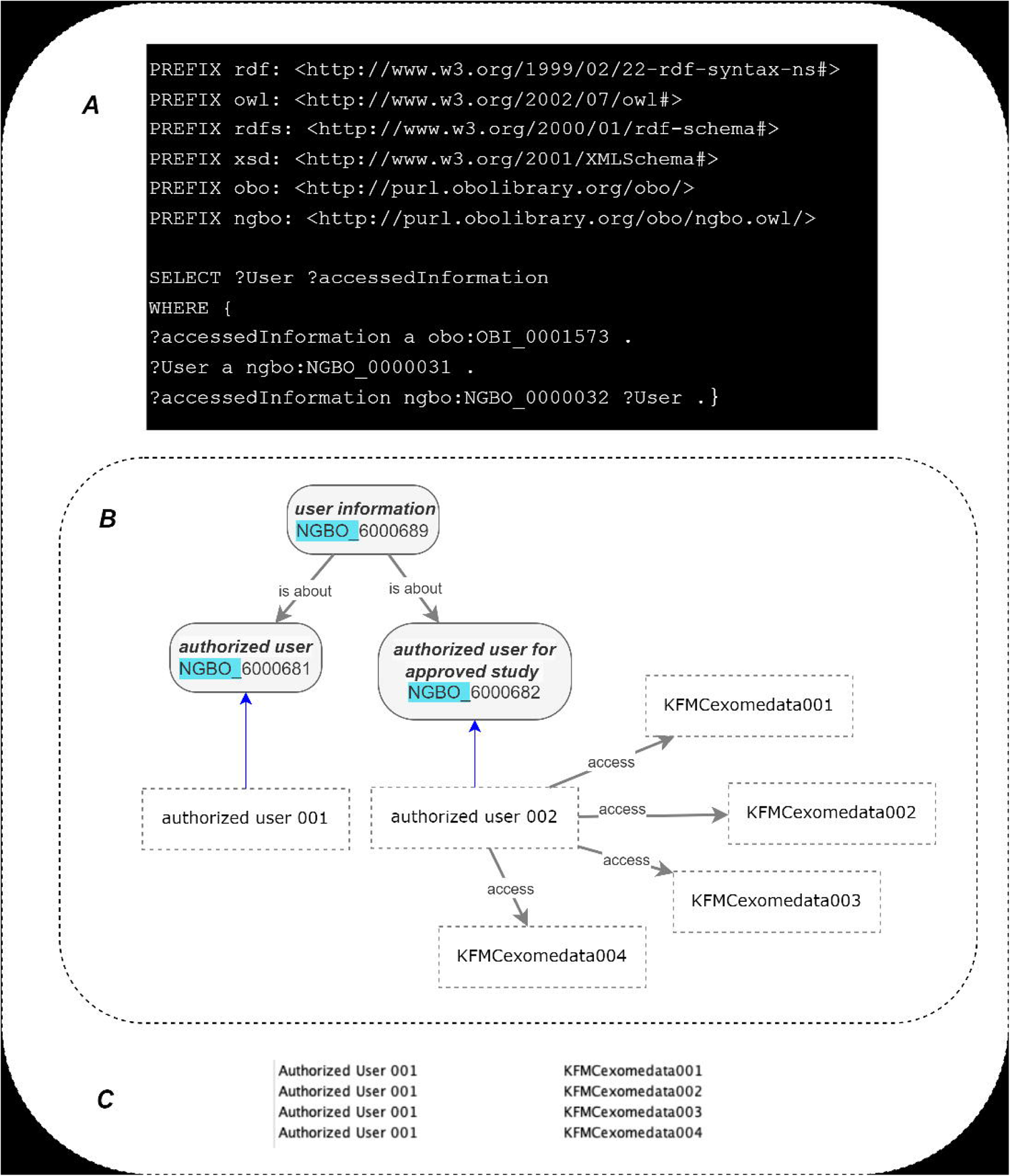
An illustration of the target region enrichment assay and assay participants.

An example of NGBO curated term is the “target enrichment region assay.” Recal that information on the target enrichment process is currently not standardized in any previous minimum information requirements or data standardization protocols. The current practice is to store the relative information as part of textual entities (computationally inaccessible). In NGBO, using the OWL class-relationship, we define axioms and formal definitions, which enable computational access to (at least some parts of) the meaning of an entity. They allow the inference of logical consequences and querying. The following formal definition, for example:

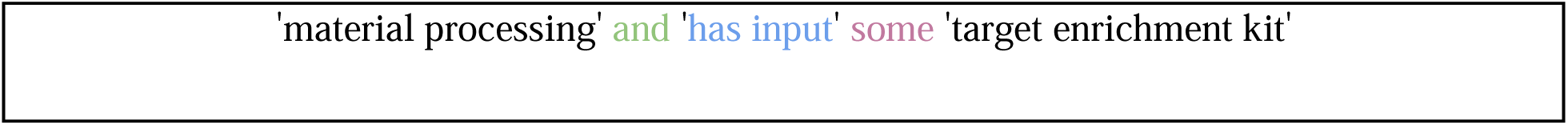

is a statement that defines part of the “target region enrichment assay” definition (See Figure 1). This is important because, in different “target region enrichment assay” methods, the sensitivity, specificity, uniformity of coverage, and reproducibility depend on the protocol [26]. More clarity regarding the enrichment process can lead to improved reuse and interpretation of the data. Here, we also followed the list of variables recommended to be reported by CAP [8]

### Ontologizing contextual data structure for wet bench analysis

Given that sequencing technologies are rapidly evolving, it is essential to accommodate traceability for each processing step in real-time [27]. The NGBO definition of the OMICS data generation assay is “a planned process that generates OMICS data through a wet bench process, such as sequencing techniques.” Each NGBO-defined entity that describes OMICS data generation assays “process” have several participants and related concepts suggested as the minimum information requirements for omics data generation assays (see supplementary file Table 2); each omics data generation assay specifies input/s and output/s to achieve its objective. For example, a “**DNA processed specimen**” **-**defined as “a processed specimen which is the output of preparing a DNA sample for further analysis”-is captured as an input of the process using the “**has_specified_input**” relation. The measurement datum “**output data file,”** defined as the “data file generated by Omics data generation assay using a measurement device” is captured as an output of the processes using “**has_specified_output**.” Here, NGBO enabled long-term archiving of both data and contextual data, and they can be made available using the standard technical procedure; in other words, both the data files generated by sequencing assay, for example, and all related contextual data are retrievable by their NGBO identifier using a standardized communications protocol (e.g., query language). Additionally, contextual data can be exchanged and used across different applications and systems as ontologised contextual data enable the integration of ontologised datasets from different sources.

### Ontologizing contextual data structure for dry bench analysis

Other NGBO development efforts involve concept curations representing bioinformatics analysis and bioinformatics software. Similar to other processes presented earlier in this paper, NGBO captures bioinformatics process participants, such as specified inputs and outputs, and information regarding process execution, including executor name, date, and run ID. NGBO also captures the plan specifications executed, including algorithms, protocols, and bioinformatics software (See Figure 2). “**bioinformatics analysis software”** is defined as software designed to extract meaningful information from the mass of molecular biology data. As part of developing NGBO, we propose a suggested minimum information requirement for bioinformatics software (see supplementary file Table 3) and minimum information requirements for SNP reporting (see supplementary file Table 4).

**Figure 2:**
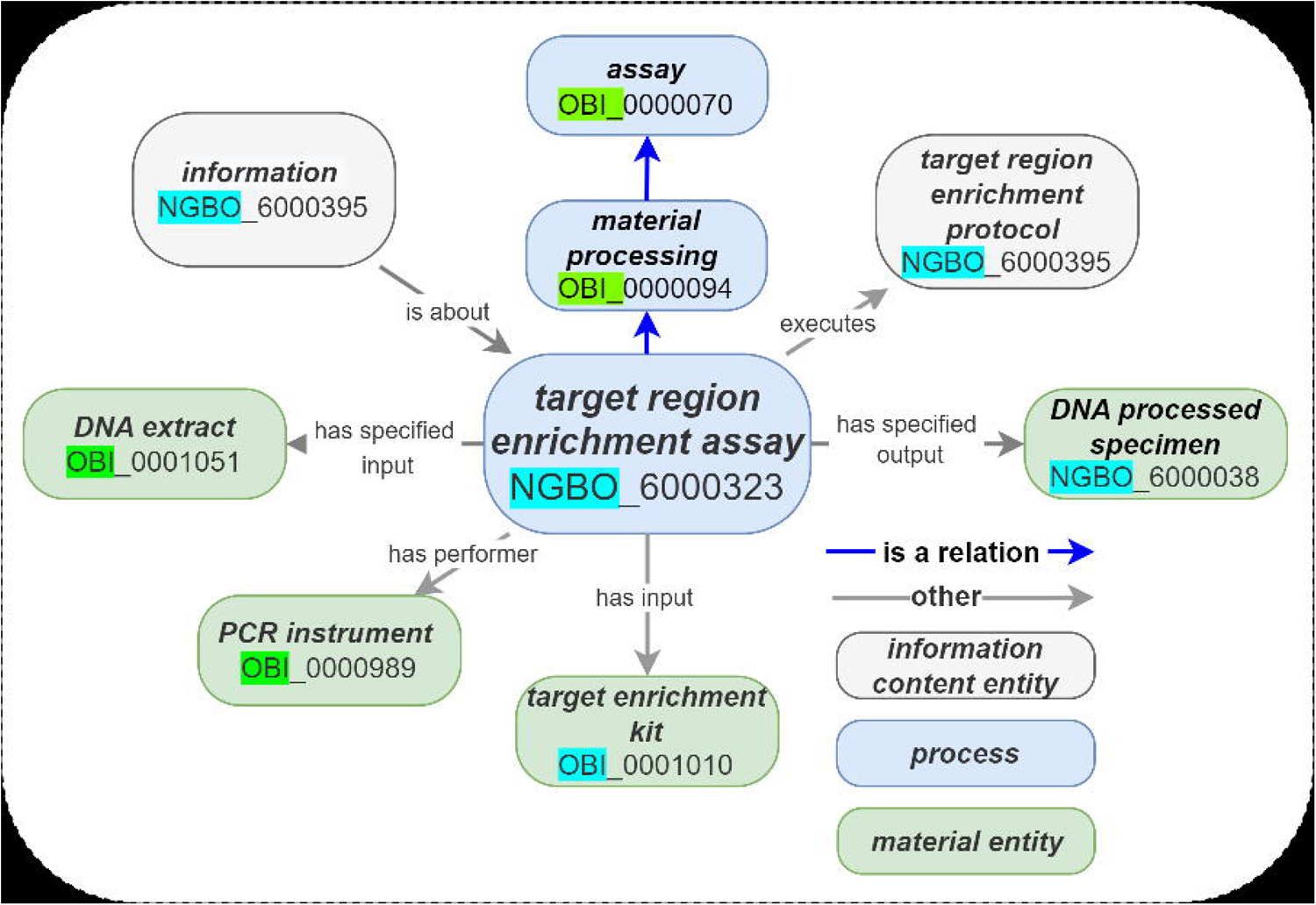
Illustration of the bioinformatics analysis and a selection of possible pipeline components.

NGBO captures bioinformatics software components, including algorithms, databases, and protocols for analyzing biological data. Additionally, the ontology model links the wet lab process with the bioinformatics analysis process to apply a bioinformatics analysis to specimen-derived data that can be linked to the original specimen (See Figure 3). The version traceability of bioinformatics analysis software or its components is captured using a “version-number” datum. The generic “**is about**” relation, as shown in the following axiom.

**Figure 3:**
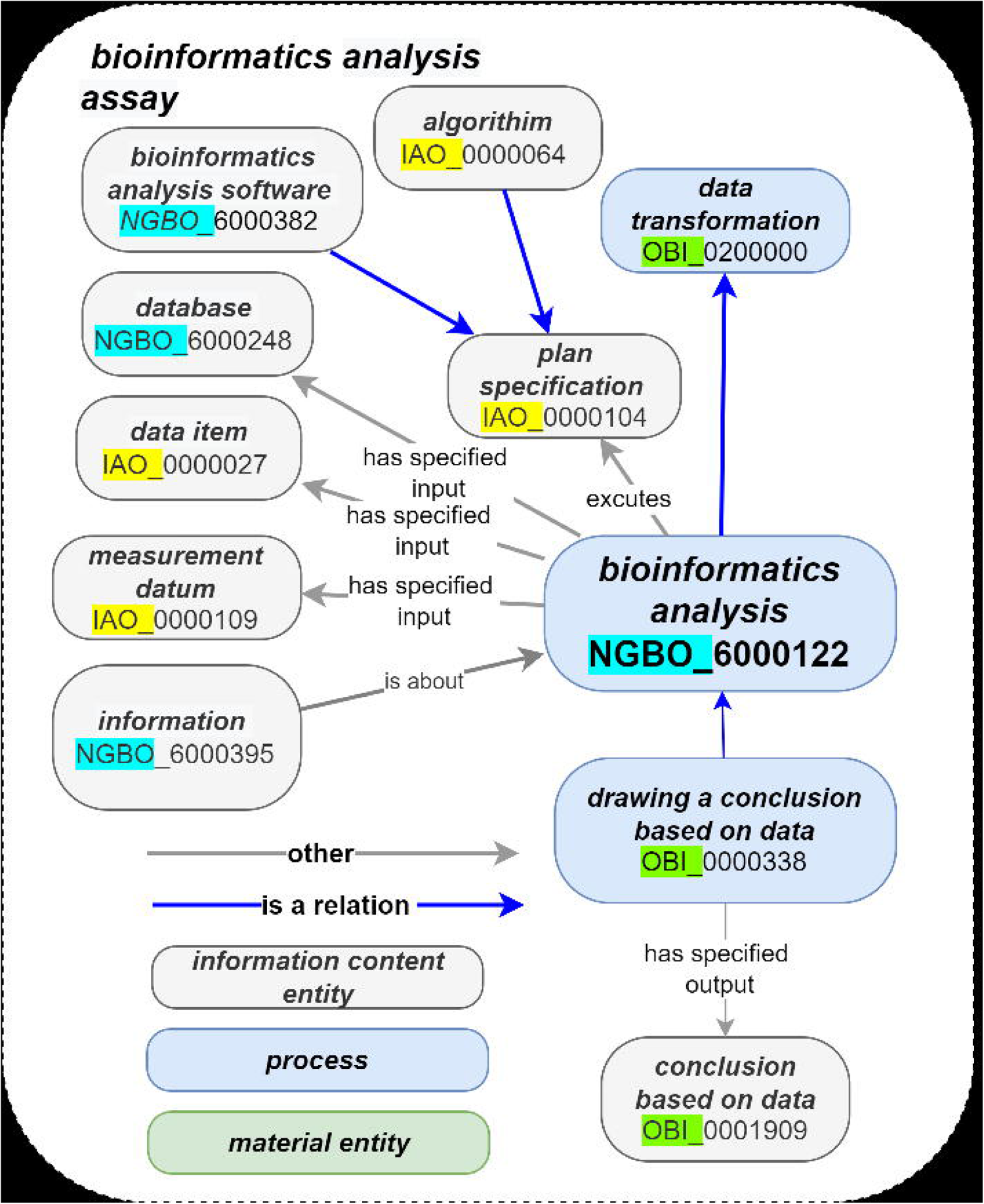
The representations of the relationship between wet lab data generation assays and bioinformatics analysis.

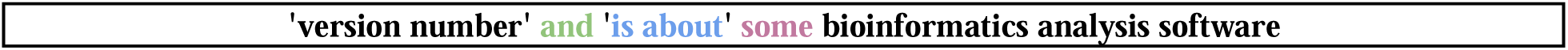

Information regarding command-line flags or parameters (when applicable) is essential for data re-analysis (if and when needed). Figure 4 illustrates an example of NGBO capturing the information using the -A INT command line flag while running the Burrows–Wheeler Aligner algorithm. NGBO captures data management activities, including de-identification, storage, and transfer. In addition, the representation includes a description of the storage data (e.g., data file URL) and protocols (e.g., data-transfer protocol).

**Figure 4:**
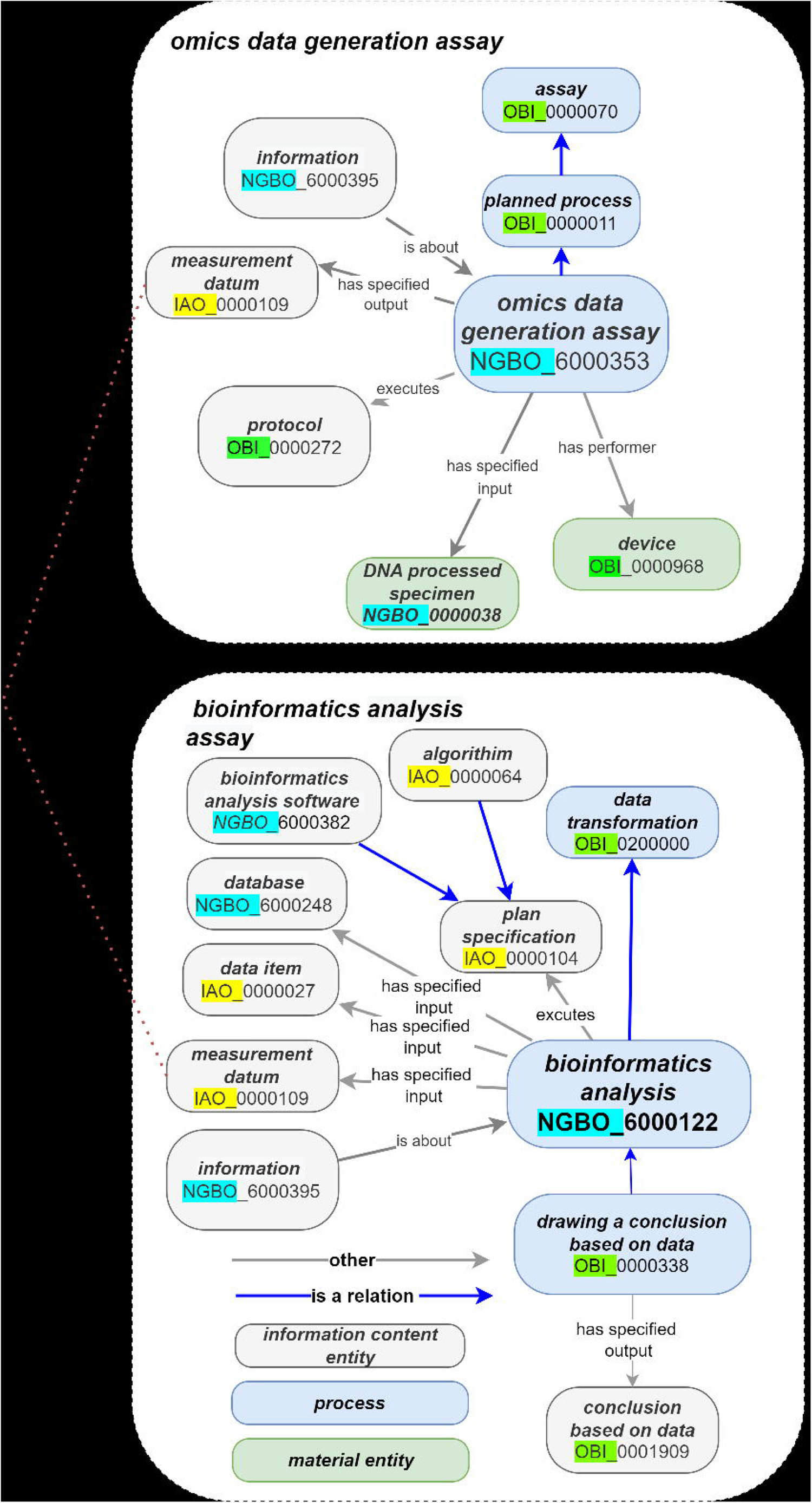
Illustration of capturing Burrows–Wheeler Aligner algorithm command line parameters.

### NGBO evaluation

#### NGBO evaluation by OBO Foundry

NGBO has been evaluated and accepted by the OBO foundry. The ontology has gone through automated checks (based on the principles below) and manual review and voting for acceptance by the OBO Foundry Operational Committee members. The submission request can be found at https://github.com/OBOFoundry/OBOFoundry.github.io/issues/1819#issuecomment-1396905637

The following summarizes how NGBO meets each of the OBO Foundry Principles:

- Open - NGBO is freely available and licensed under CC BY 4.0 on GitHub and OLS
- Common Format - NGBO is available in a common formal language (XML/rdf and owl) in concrete syntax.
- URI/Identifier Space -NGBO has a unique IRI in the form of an OBO Foundry permanent URL (PURL); http://purl.obolibrary.org/ngbo
- Versioning - The latest version of the ontology can be found at https://github.com/Dalalghamdi/NGBO/blob/master/ngbo.owl; details on previous versions can be found on (https://github.com/Dalalghamdi/NGBO).
- Scope - This ontology includes real-world physical objects such as specimens, computerized objects such as datasets, and planned processes and related specifications.
- Textual Definitions - The ontology has textual definitions for all its classes
- Relations - Relations are reused from the Relations Ontology (RO), except for four relations defined by NGBO.
- Documented Plurality of Users - The intended end-users are biobanking organizations, data collectors, omics researchers, and decision-makers.
- Commitment To Collaboration - new terms introduced by NGBO will be submitted to appropriate OBO foundry domain ontology when possible. NGBO also accepts and reviews requests from the community to edit/add terms through issue requests here (https://github.com/Dalalghamdi/NGBO/issues)
- Locus of Authority - The primary contact “Dalia Alghamdi” is responsible for communications between the community and NGBO developers.
- Maintenance - The ontology needs to reflect changes in scientific consensus to remain accurate over time.
- Documented plurality of users - The SHGP is a Saudi national project initiated in 2014.

On top of the program’s main objectives is to build a genetic database for the Saudi population and enable scientists and researchers to benefit from the genetic information in the project. Typically, like many similar projects worldwide, the contextual data required by researchers for data discovery and reuse are intermixed in separate silos, including bioinformatics software information, laboratory SOPs, and reports. On the one hand, while the growing data is being distributed between silos, the number of potential users is also growing. As a result, it is challenging to constantly manage and govern this data while making data universally discoverable to potential consumers. On the other hand, contextual data fields stored in different locations might lead to inconsistencies, inhibit a complete view of entities, and need more data integrity. Here, the implementation of NGBO enables the breaking down of walls between different pieces of information in one place. NGBO covers terms specific to wet and dry bench processes and provides a framework that explains how NGBO and other ontology terms relate to each other in the context of an experiment.

#### NGBO evaluation by CQs and reasoners

To validate and test the adequacy of the ontology, we referred to the method used in a previous study [15]. Description logic (DL) queries over an OWL file populated with MoC data instances and false positive and negative entries at the instance level. The MoC data were imported into the NGBO OWL file using the Celfie plugin [28]. The DL queries are implementations of the previously proposed CQs.

When evaluating the quality of NGBO, we considered two principles:

1. Clarity: ontology does not create any misinterpretation during query execution or reasoning.
2. Consistency: all individuals were accurately categorized when the reasoning procedure was completed.

Reasoning is deriving implicit facts from a set of given explicit facts. In NGBO, these facts are expressed in OWL and stored in RDF triplestores. For example, the following fact: “a bioinformatics analysis executes an algorithm” can be expressed in an ontology at the class level as “ ‘bioinformatics analysis’ [<u>NGBO:6000122</u>] executes some algorithm [<u>IAO:0000064</u>]“, while an instance of this fact involving a particular job run and software reference: “bioinformatics analysis X123 executes algorithm Y456” can be stored in a triplestore. A reasoner is a software application that uses an ontology to infer relationships that should hold between existing classes and their instances. This allows them to categorize entities under ontological hierarchies, and it allows them to identify contradictions. To validate whether or not NGBO is consistent, we used HermiT [29] and Pellet’s [30] reasoners. And to validate the ontology ability to answer the CQs, we used description logic queries available in (See Supplementry file CQ). To check NGBO’s ability to answer the questions.

### Semantic web application for organizing and federated querying of biobank data

Data describing biobank resources often contain unstructured text information with insufficient framework for computational processes, including automated data discovery and integration. As a proof of concept, we demonstrate how semantic web technologies (e.g., SPARQL, RDF, ontologies) can be used to build an application to improve querying and integrate biobank specimens and specimen-derivative data, including genomic sequences. We set up a web application [31] for end-users with appropriate functions to import, query, and retrieve data. The semantic web application is freely available and licensed under Apache License 2.0.

## ADDITIONAL FEATURES

### Specimen tracking and traceability

The detailed documentation of all specimen processing activities is critical for many reasons, including accreditation and regulatory body requirements, collaboration and data reproducibility, and audibility. For example, CAP requires traceability of all standard operating protocols for DNA/RNA specimen preparation, library preparation, sequence generation, and bioinformatics analysis.

NGBO includes terms describing specimen processing, material entities, and information entities, including information about designated team members or personnel who perform any specimen- or data-related activities. Additionally, a set of relations established between individuals was used to identify directly or indirectly affected processed specimens and datasets. From within NGBO relations (object properties), two major relations are used for tracking and traceability purposes: “trace from” and “track to.” The first relation, **“trace from,”** includes sub-relations that enable top-down tracing, Sub-relations of the relation **“trace from”** are: **“has performer,” “executes,” “has specified input,” “has specified output,” and “part of,” “trace from”** enables bottom-up tracing. On the opposite side, **“track to”** and its sub-relations, the inverse properties of **“trace from,”** allows a top-down tracking of activities. Here, when the relations between individuals of classes are described, for example, **“has performer,”** the relation “trace_from” can be inferred via ontology reasoning. The “trace from” and “track to” relationships can be examined in Semantic Query-Enhanced Web Rule Language (SQWRL) requests for bottom-up or top-down tracing; an example is given in Figure 5.

**Figure 5:**
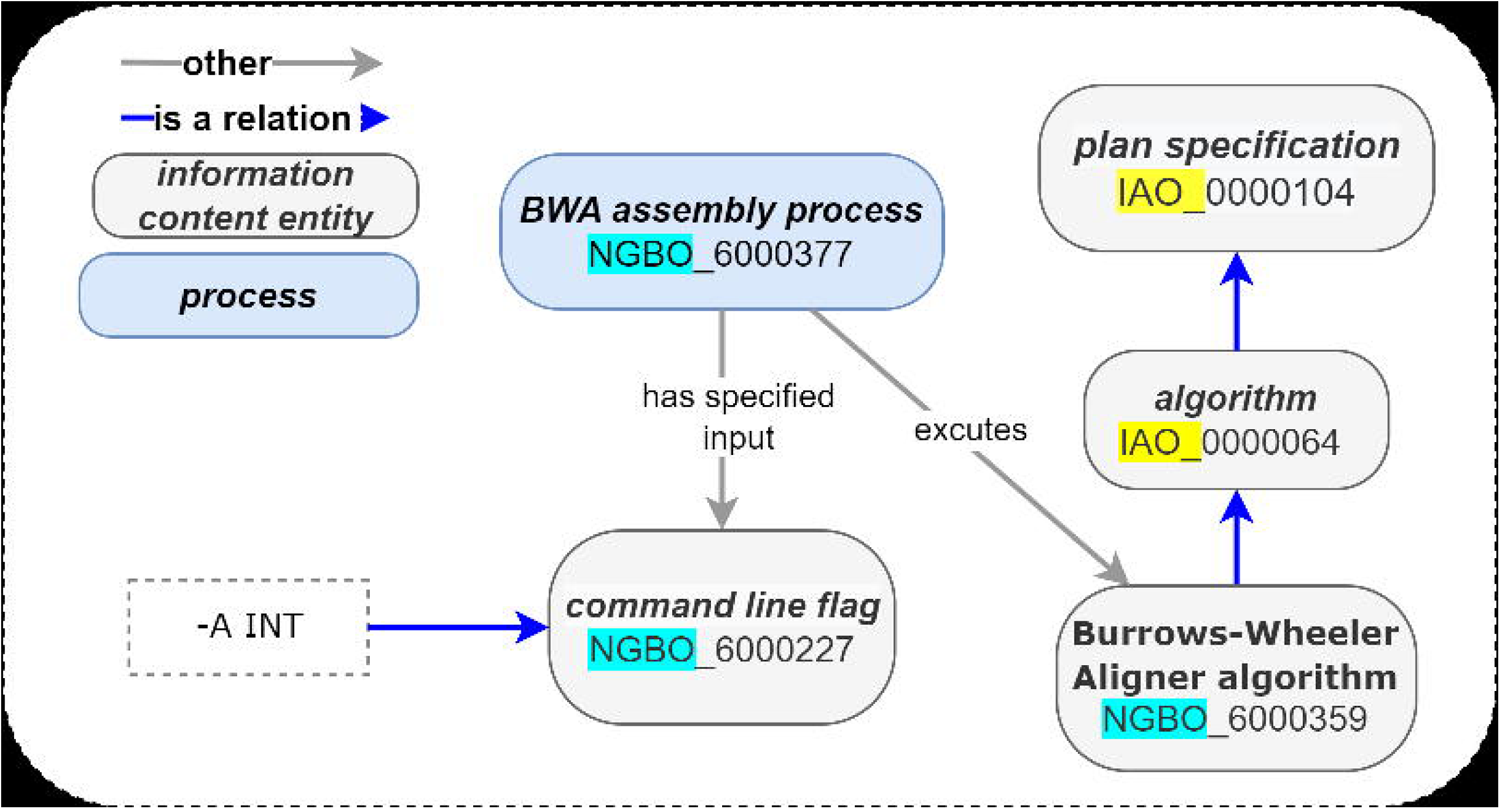
Using NGBO for specimen tracking and traceability. (A) An example of relations between individuals. (B) A top-down tracing SQWRL query using the “trace_from” relationship. (C) Table of query results. The traceability results of the query are derived through an ontology reasoning system and selected in variable tracing. The variable “ExamAssay” contains instances linked through the “trace_from” relationship.

### Data privacy and user access control

When dealing with biobanking data, organizations must ensure justified data access for the right person. In this work, we depended on graph-based representations of contextual data based on relations links between data. We propose four different classes (“data owner,” “guest,” “authorized user,” and “authorized user for approved study”) that enable the exploitation of semantic relations between nodes using the “accessed_by” object property for data access that allows for meaningful access protocols. In traditional databases, applications must check the permission of a specific material with every run; however, graph queries can efficiently follow network connections through relationships and check the permission of a specific material with every run Figure 6 shows an example of a SPARQL query used to find all users accessing exome data files.

**Figure 6:**
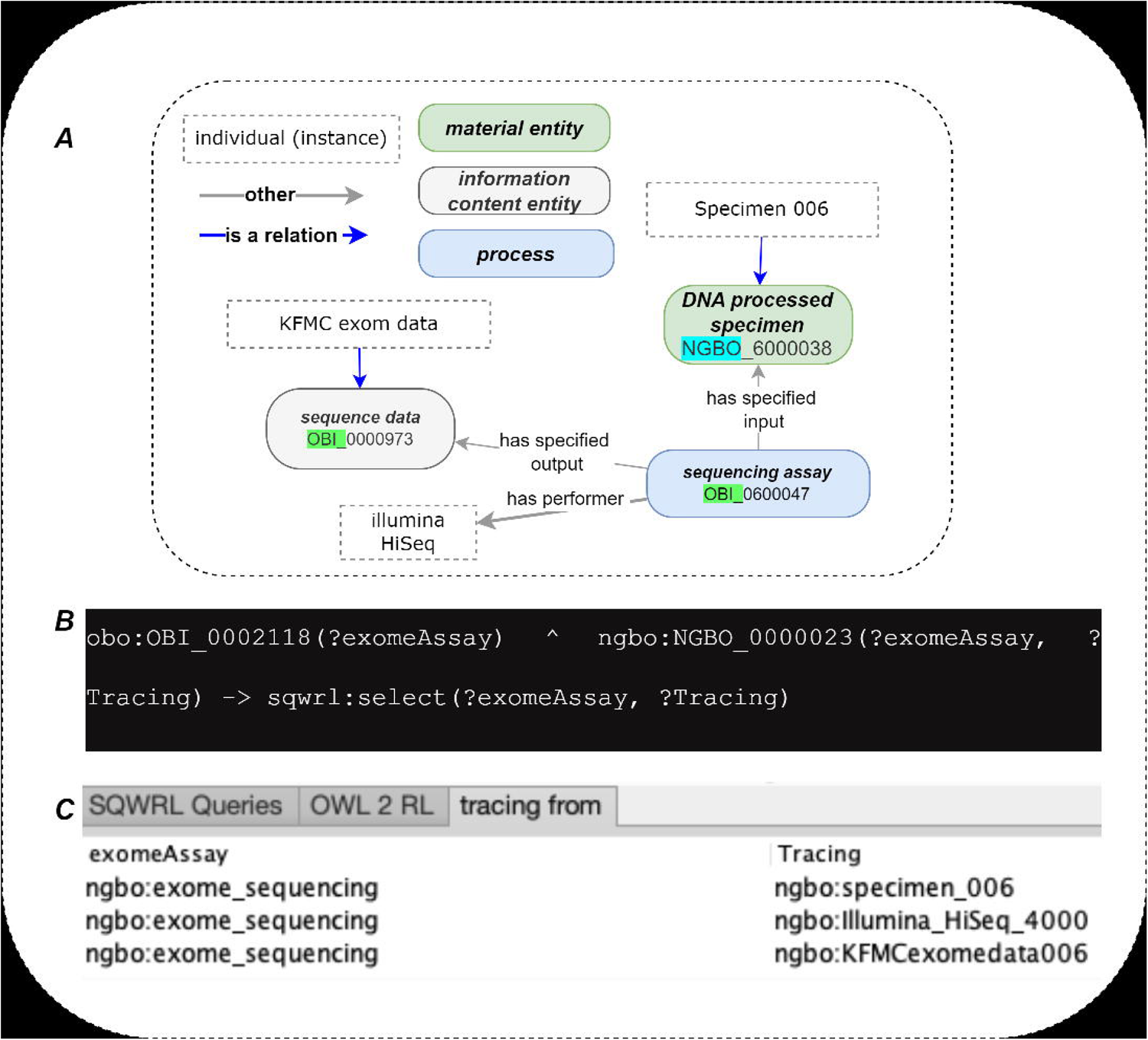
Using NGBO for data privacy and user access controls. (A) A SPARQL query to find who can access a set of exome files. (B) An example of relations between individuals. (C) SPARQL query result, all individuals were linked to the set of exam data files through the relation “access_by” selected in the variable?User. Here, “authrised user 002” has access to the approved data sets through “access”, while “Authrised user 001” cant access the same dataset as there no relationship established between the user and the dataset.

## DISCUSSION

Biobanks bring together specimen donors, their specimens, associated data, and the researchers interested in utilizing the information [32]. In this paper, we present the development of NGBO, a biobanking application ontology that applies semantic annotations to contextual data obtained from omics experiments. NGBO fills the need for semantically enabling the discovery and integration of omics datasets and realization of FAIR data representation, which will impact the efficiency of finding, integrating, and re-using biobanking data of interest. NGBO development followed established principles, standards, and resources, including the OBO Foundry principles. Furthermore, we captured information from sources, including the College of American Pathologists. This scope of NGBO includes analytical wet bench assays, bioinformatics analysis, and data management processes. Integrating ontology directly into biobanking records will enable broad-range data sharing that links participants, biobanks, and researchers. Additionally, with the increasing number of genome and exome sequencing, there is a potential to use ontology-based algorithms to integrate and implement the results from basic biology research into clinical decision-making [33].

The ontology is maintained in GitHub and will continually improve based on community input and user feedback. Since NGBO is an application ontology, NGBO-defined terms will be submitted to relevant domain ontologies. In the future, we will continue the data curation effort by completing the metadata management proposal for the SHGP and King Fahad Medical City metadata management proposal. We will also consider other sources pointed out by the reviewers of NGBO, starting with the NCBI bio-sample [34].

## Supporting information

(see supplementary file Table 4).

(see supplementary file Table 2)

(see supplementary file Table 3)

## Acknowledgments

The authors wish to acknowledge the Saudi Diagnostics Lab (SDL) members for their helpful discussions and support. In addition, this work is funded by King Fahad Medical City (KFMC), Saudi Arabia.

