## Supplementary material for "NGBO: Introducing -omics metadata to biobanking ontology": (see supplementary file Table 4).

| **Data item (Interface label)** | **Ontology label** | **Definition** | **Guidance** | **Example of use** |
| --- | --- | --- | --- | --- |
| Detected gene mutation | detected gene mutation  (NGBO) | A conclusion based on data about a permanent change in the DNA sequence that makes up a gene  *(This entity is only used for data structuring purposes)* | Select value from the pick list. If not available, request a new term by creating a new issue here: https://github.com/Dalalghamdi/NGBO/issues | **KRAS mutation** |
| Gene names by HUGO Gene Nomenclature Committee (HGNC) | HUGO gene nomenclature committee gene name  (NGBO) | The biological database by HGNC contains approved unique symbols and names for human loci, including protein-coding genes, RNA genes, and pseudogenes. HUGO Gene Nomenclature allows clear scientific communication. | Select value from the pick list. If not available, request a new term by creating a new issue here: https://github.com/Dalalghamdi/NGBO/issues | **MTHFR** |
| Genomic coordinates | Reference genome coordinate  (NGBO) | Reference genomic coordinate is a data item that describes the start and end position of a gene (or genetic element) on a chromosome. | Enter Genomic coordinates, and include the chromosome name, start position, and end position in the following format: chr1:1234570-1234870 | **chr1:1234678-1234567.** |
| RefSeq accession (transcript reference number) | RefSeq accession identifier  (GenEpio) | RefSeq accession is a data item describing the set of character sequences of a RefSeq database entry. | Enter RefSeq record, a distinct accession number format that begins with two characters followed by an underscore (e.g., NP_) | **nm_00739.1** |
| Reference genome name and version | reference genome version number  (NGBO) | reference genome datum which represent reference genome edition | Select value from the pick list. If not available, request a new term by creating a new issue here: https://github.com/Dalalghamdi/NGBO/issues | **GRCh37** |
| Coding information | nucleic acid substitution  (NGBO) | A data item about the biological process in which one nucleic acid is replaced by a different nucleic acid. | Enter replacement information of one nucleotide in a DNA with a different nucleotide in the following format:  **c.973C>T** | **c.973C>T** |
| Variant allele frequency (VAF) | allele frequency  (STATO) | Variant allele frequency is a data item that describes the percentage of sequence reads observed matching a specific DNA variant divided by the overall coverage at that locus. VAF determines somatic or germline mutations. | Enter the percentage of sequence reads observed matching a specific DNA. | **Usually shown as a percentage (%)**  **60%** |
| Amino acid replacement data item | amino acid substitution  (NGBO) | A data item about the biological process in which one amino acid is replaced by a different amino acid. | Enter replacement information of one amino acid in a protein with a different amino acid in the following format:  p. Trp26Cys | **p.Arg325Cys** |

Table 4: Suggested Minimum information requirements for SNP reporting
