## Supplementary material for "NGBO: Introducing -omics metadata to biobanking ontology": (see supplementary file Table 2)

| **Data item (Interface label)** | **Ontology label** | **Definition** | **Guidance** | **Example of use** |
| --- | --- | --- | --- | --- |
| Sequence run executer name | Name of data generation assay executor  (NGBO) | Name of personnel who executed the sequencing run | Enter the name of personnel who executed the sequencing run | Dalia Algahmdi |
| Sequence run execution date | Sequencing run date  (GenEpio) | Date the sequencing run was performed | Enter the date the sequencing run was performed. The required granularity includes year, month, and day. | 21/05/2022 |
| Sequencing run execution note | Assay execution note  (NGBO) | A textual entity of relevant notes or instructions regarding data generation assay execution | Enter free text of relevant notes or instructions regarding your sequence run. Do not enter any information describing entities mentioned as part of  this table (e.g. Name of data generation assay executor or Target gene specification) |  |
| Any protocols used in data generation assay | Protocol  (OBI) | A plan specification that has sufficient level of detail and quantitative information to communicate between investigation agents, so that different investigation agents will reliably and independently be able to reproduce the process | Select value from the pick list. If not available, request a new term by creating a new issue here: https://github.com/Dalalghamdi/NGBO/issues | Target enrichment protocol (NGBO)    Adapter ligation protocol (NGBO) |
| Information about sequencing target region | Target gene specification  (OBI) | A directive information specifying a coding genomic region, which is the focus of a planned process such as an assay in an environmental gene survey | Enter gene panel name, users must clearly clarify if the gene panel is an in-house gene panel or public gene panel. | Autism gene panel |
| Assay kits used in data generation assay | Assay kit name  (NGBO) | Processed material that contains a set of articles or equipment needed for a specific purpose | Select value from the pick list. If not available, request a new term by creating a new issue here: https://github.com/Dalalghamdi/NGBO/issues | QIAseq Human Exome Kit |
| Versions of assay kits | Assay kit version  (NGBO)  Sequencing kit version  (GenEpio) | Version of assay kit | Select value from the pick list. If not available, request a new term by creating a new issue here: https://github.com/Dalalghamdi/NGBO/issues | User entry: 1.0 |
| Any devices used in data generation assay | Device  (OBI) | A material entity that is designed to perform a function in a scientific investigation, but is not a reagent | Select value from the pick list. If not available, request a new term by creating a new issue here: https://github.com/Dalalghamdi/NGBO/issues | Illumina HiSeq 4000 |
| Specimen identifiers | Identifier  (ISO) | An identifier is a label that specifically refers to (identifies) an entity (instance/type). | Enter identifier of processed specimen, which is the output of preparing a DNA sample for further analysis | Specimen28728 |
| Omics data generation assay output data file | Data file name  (NGBO) | Data file generated by Omics data generation assay using a measurement device | Enter data file name | Read27828.fasta |
| output data file format | data file format  (NGBO) | Data file format is a standard way that information is encoded for storage in a computer file | Select value from the pick list. If not available, request a new term by creating a new issue here: https://github.com/Dalalghamdi/NGBO/issues | FASTA |

Table 2. Suggested minimum information requirements for omics data generation assay.
