## Supplementary material for "NGBO: Introducing -omics metadata to biobanking ontology": (see supplementary file Table 3)

| **Data item (Interface label)** | **Ontology label** | **Definition** | **Guidance** | **Example of use** |
| --- | --- | --- | --- | --- |
| Bioinformatics analysis execution date * | bioinformatics analysis execution date  (NGBO) | Date on which analysis was performed | Enter the date the bioinformatics analysis was performed. The required granularity includes year, month, and day. | 21/05/2022 |
| Bioinformatics software executor name * | name of bioinformatics analysis executor  (NGBO) | Name of personnel who executed the analysis. | Enter the name of personnel who executed the bioinformatics analysis (text). | Dalia Algahmdi |
| software run ID * | software run identifier  (NGBO) | An identifier that denotes the unique run ID of the code executed | Entre software run ID | PID: 346524 |
| Bioinformatics analysis execution note* | assay execution note  (NGBO) | A textual entity of relevant notes or instructions regarding bioinformatics analysis execution. | Enter free text of relevant notes or instructions regarding your sequence run. Do not enter any information describing entities mentioned as part of this table (e.g. software run ID, Bioinformatics software version number) | This analysis did NOT identify genetic variants that may be responsible for other diseases unrelated to this individual’s clinical presentation. |
| Bioinformatics software name* | software name  (NGBO) | A name of the Bioinformatics software utilized. | Select value from the pick list. If not available, request a new term by creating a new issue here: https://github.com/Dalalghamdi/NGBO/issues | 1) JWES pipeline.  2) CLC Genomics Workbench. |
| Bioinformatics software version number * | software version number  (NGBO) | A version number of bioinformatics analysis software. | Entre software version number, if the software/pipeline is an in-house product, it is recommended to follow numbering system i.e. 3.01 or date-based following [ISO-8601](https://www.iso.org/iso-8601-date-and-time-format.html), ie. date format “YYYY-MM-DD” | v. 3.01 |
| Bioinformatics software description | software description  (NGBO) | A description of the Bioinformatics bioinformatics software. | Do not enter any information describing entities mentioned as part of this table (e.g. software run ID, Bioinformatics software version number) | QIAGEN CLC Main Workbench is used by tens of thousands of researchers all over the world for DNA, RNA, and protein sequence data analysis.. |
| Bioinformatics software Documentation URI | software Documentation URI  (NGBO) | A URI of associated manuals, documents, or guidance | Entre a URI for a written text or illustration that accompanies computer software or is embedded in the source code | https://github.com/drzeeshanahmed/JWES-Variant |
| Input data file name | Data file name  (NGBO) | Data file generated by Omics data generation assay using a measurement device | Enter data file name | Read27828.fasta |
| Input data file path* | data file URL (NGBO) | A URL at which the data file file is stored. | Enter data file URL | e.g. \\server01\kfmc\path |
| Data file format* | data file format  (NGBO) | Data file format is a standard way that information is encoded for storage in a computer file | Select value from the pick list. If not available, request a new term by creating a new issue here: https://github.com/Dalalghamdi/NGBO/issues | FASTA |
| Reference genome download URI | reference genome URI (NGBO) | A reference genome datum that represents a location where reference genome can be accessed online | Enter reference genome download URI | https://www.ncbi.nlm.nih.gov/genome/guide/human/ |
| Reference genome build* | reference genome identifier  (GenEpio) | Identifier for a reference genome (used to build or compare an assembly) that is used in the genome browsers and in the community | Entre reference genome symbol. | GRCh38 |
| Bioinformatics software parameter name* | software parameter name  (NGBO) | Name of a software option | Enter bioinformatics software parameter name, do not add any extra characters. | -q |
| Bioinformatics software parameter value* | software parameter value  (NGBO) | The value to be fed into software option | Enter bioinformatics software parameter value, do not add any extra characters. | INT |
| Command-line flag* | command line flag  (NGBO) | A symbol that is used to specify options for command-line programs. | If applicable, enter any Command-line flag used. | Appropriate command-line flags for running bioinformatics pipeline, for example, bwa - Burrows-–Wheeler Alignment l |
| Bioinformatics analysis output data file URI * | data file URL (NGBO) | A URL at which the data file file is stored. | Enter data file URL | e.g. \\server01\kfmc\path |

Table 3. Suggested minimum information requirements for bioinformatics software. * (asterisk) indicate mandatory requirement when applicable
