## Supplementry file CQ) for "NGBO: Introducing -omics metadata to biobanking ontology"

**A.1 Supplementary method - An examples of description logic queries expressing competency questions.**

Search for disease **x (adenocarcinoma specimen)** omics-derived data **(DNA sequence data)** across all available repositories.

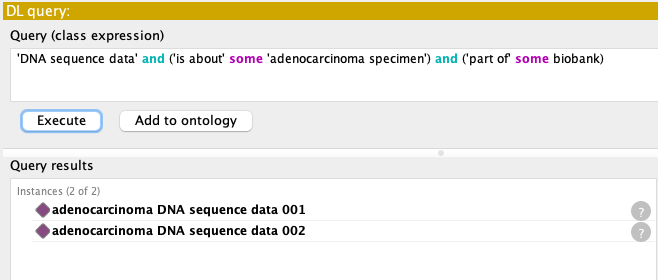

Search for disease **x (adenocarcinoma specimen)** omics-derived data **y (DNA sequence data)**  available in repositories **a, b (King Fahad Medical City biobank & King Faisal Specilist Hospital)**  with a mutation in gene **z (KRAS mutation)**.

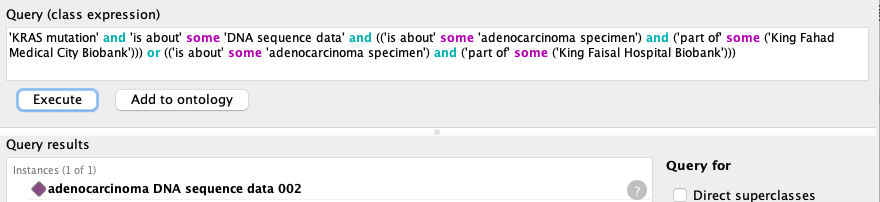

Data format **x (fasta files)** for omics-derived data **y (DNA sequencing data)** for disease **z (autism)** filtered by life stage **(pediatric patients)** available in repositories **a (King Fahad Medical City)**

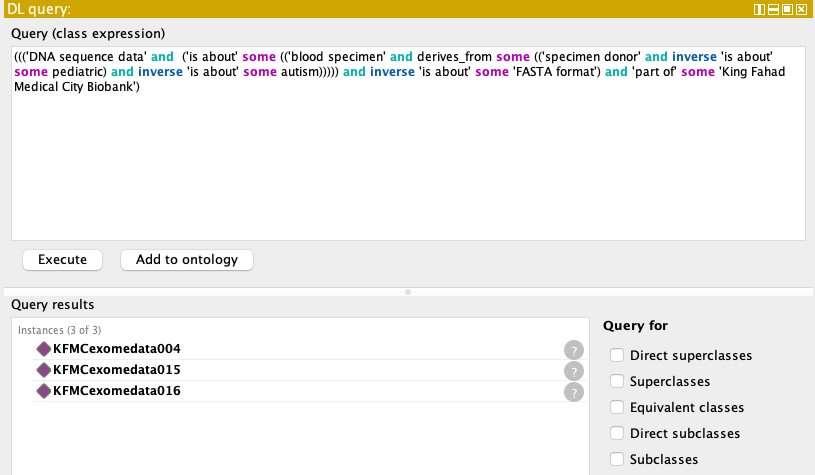

Number of data formats **x (fasta files)** for omics-derived data **y (DNA sequencing data)** for disease **z (autism)** filter by life stage **(pediatric patients)** available in repositories **a (King Fahad Medical City)**

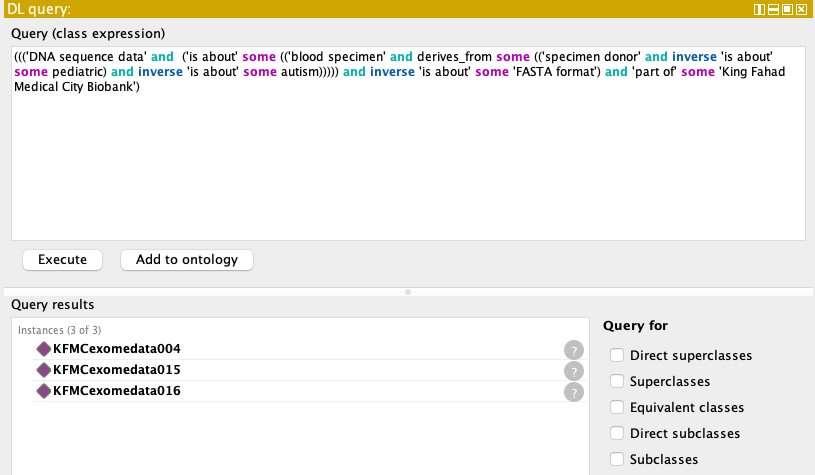

Search for omics-derived data for disease **x (bladder cancer)** and subset it by smoking history

**(current reformed smoker for less than 15 years history), (current reformed smoker for more than 15 years history).**

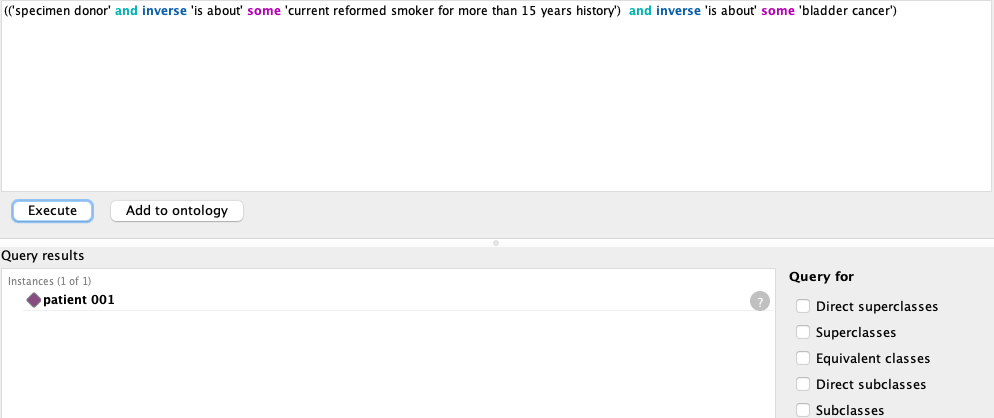

Search for omics-derived data **(DNA sequencing data)** **x** with mutation **y (FGFR3)** and **z (PIK3CA)**

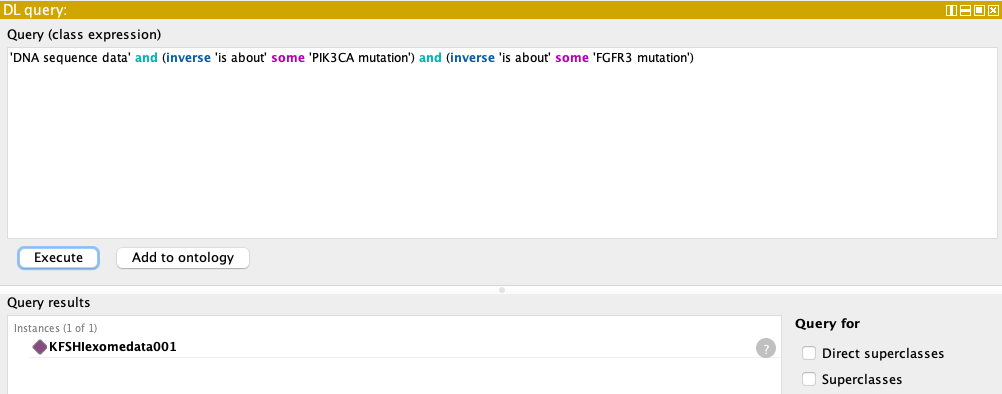
